# A disinhibition-based model of CA1 place field sequences

**DOI:** 10.64898/2026.10.08.757765

**Authors:** Laura Green, Mihály Vöröslakos, György Buzsáki, John Rinzel

## Abstract

Because of the paucity of pattern-generating recurrent excitation in the CA1 hippocampal region, place field sequences are assumed to be inherited from upstream regions. Alternatively, disinhibition may play a role in shaping pyramidal cell activity. We propose a model of CA1 activity using one layer of excitatory cells and one layer of inhibitory cells, with local excitatory-to-inhibitory connectivity, broad inhibitory-to-excitatory connectivity without recurrent excitatory connections, and broad inhibitory-inhibitory connections. The layers need not follow a topographic organization; instead they can represent sequences of place cells structured by local inhibition. Our mathematical model of CA1 neurons with data-inspired connectivity demonstrates place fields sharpened by disinhibition, mimicking a recently discovered decrease in inhibitory input within place fields and matching in vivo patterns of disinhibition. Mathematical analysis of our model suggests the importance of inhibitory-to-excitatory connections in control of place cell firing rate and sparsity. We also demonstrate that a disinhibition-based mechanism of pattern generation is more flexible without recurrent excitation than the recurrent excitation-rich CA3-like model. Finally, we demonstrate disinhibition as an efficient mechanism for internally-generated sequential activity, where the speed of sequential change depends on both input strength and inhibitory-to-excitatory connection strength. Overall, our model suggests that theta time-scale disinhibition is a foundational mechanism of self-organized sequential activity in the CA1 region, and, possibly, in other networks.

**Author Summary:** The hippocampus is thought to be important for learning and memory. The main output of the hippocampus is through the CA1 region, which contains cells that demonstrate spatially-informative firing patterns. Previous models of place cell activity include recurrent excitatory or increases of inhibitory input within a place field, though it is well known that CA1 exhibits only weak connections between excitatory cells. We provide a population model of CA1 place cell activity without recurrent excitation that replicates many characteristics of place cell activity, such as a decrease in inhibitory input within a place field and increasingly sparse activity with familiarity in the environment. We also demonstrate intrinsic sequential activity through disinhibition. We propose that disinhibition dominates theta-timescale activity in the CA1 region of the hippocampus.

## Introduction

The hippocampus is a layered structure involved in spatial navigation, action planning and memory(1–4). The CA1-subicular regions are the main output regions of the hippocampus, projecting back to cortical areas and many subcortical/thalamic regions (5,6). The CA1 region of the hippocampus has limited recurrent excitatory connectivity, tacitly implying that the CA1 region simply relays information from CA3, where strong recurrent excitation allows for more attractor type computations such as place cell activity (7–11). However, several observations suggest CA1 performs local circuit computations: ability to perform pattern separation (12), place fields remap at different rates (13–15), and place fields and neuronal sequences persist in CA1 even without input from CA3 or the entorhinal cortex (3,16–19).

Recent work demonstrates disinhibition (20–22) and lateral inhibition within CA1 circuits (23), along with bodies of work demonstrating complex circuitry within CA1 (24,25). Canonical models of the hippocampus, from the Hopfield model (26) to more recent models involving external cues combined with internal path integration, rely on local recurrent excitation combined with broad lateral inhibition to produce sustained, localized activity (9,21,27–31). This motif of local recurrent excitation combined with broad lateral inhibition (Mexican-hat connectivity) allows for self-sustained activity that remains localized within a network. Additionally, this type of network may lead to attractor dynamics, in which a circuit is able to maintain stimulus-relevant persistent activity when the stimulus is removed. Recurrent excitation among pyramidal neurons, such as the recurrent CA3 circuit, is the assumed substrate of sequential firing (29,32,33). However, recent experiments demonstrate sequential place field activity in the CA1 region, where recurrent excitation is very sparse, even when the upstream CA3 and entorhinal inputs are inactivated (16). Thus, mechanisms other the recurrent excitation among pyramidal neurons may sustain sequential neuronal firing. Disinhibition is an alternative solution to sustained activity (21,34), potentially providing a mechanism for attractor-like activity in CA1. Other hippocampal place field models predict no change or an increase in inhibitory input within a place field and include recurrent excitatory connections (21,35–40).

In this study, we show that a disinhibition-based circuit without recurrent excitation can decrease inhibitory input within a place field during navigation. Further, this decrease in inhibition was associated with attractor dynamics, enabling the generation of sequential firing. We propose that CA1 demonstrates attractor activity during each theta cycle when place cells are active during the late-to-mid phase of theta, which is quenched during the earlier phases of theta when interneurons are most active. This leads to a theta-mediated two-phase alternation between a low-activity stable state and an attractor-like state. The result is a mechanism for improving signal-to-noise in CA1 by disinhibiting place-cell activity during each theta cycle, reducing out-of-field spikes, and maintaining the flexibility to switch between separate place-cell representations across theta cycles.

## Results

### Disinhibitory circuit motif in a spiking model decreases inhibitory input to place cells within a place field

We found that we could reproduce a decrease in presynaptic inhibitory cell activity by including local disinhibitory circuit motifs in the model, combined with broad lateral inhibition. Excitatory cells were connected with on-center connectivity to interneurons, while inhibitory cells were connected with off-center connectivity to excitatory cells. Inhibitory cells also connected sparsely to other inhibitory cells, thus completing the disinhibitory loop (Fig. 1A-C). Excitatory cells excited inhibitory interneurons that inhibited those which inhibited them, creating the feedback loop that underlies disinhibition. The same circuitry ensured that excitatory cells directed feedforward inhibition towards other excitatory cells. There were no connections among excitatory neurons.

**Figure 1.**
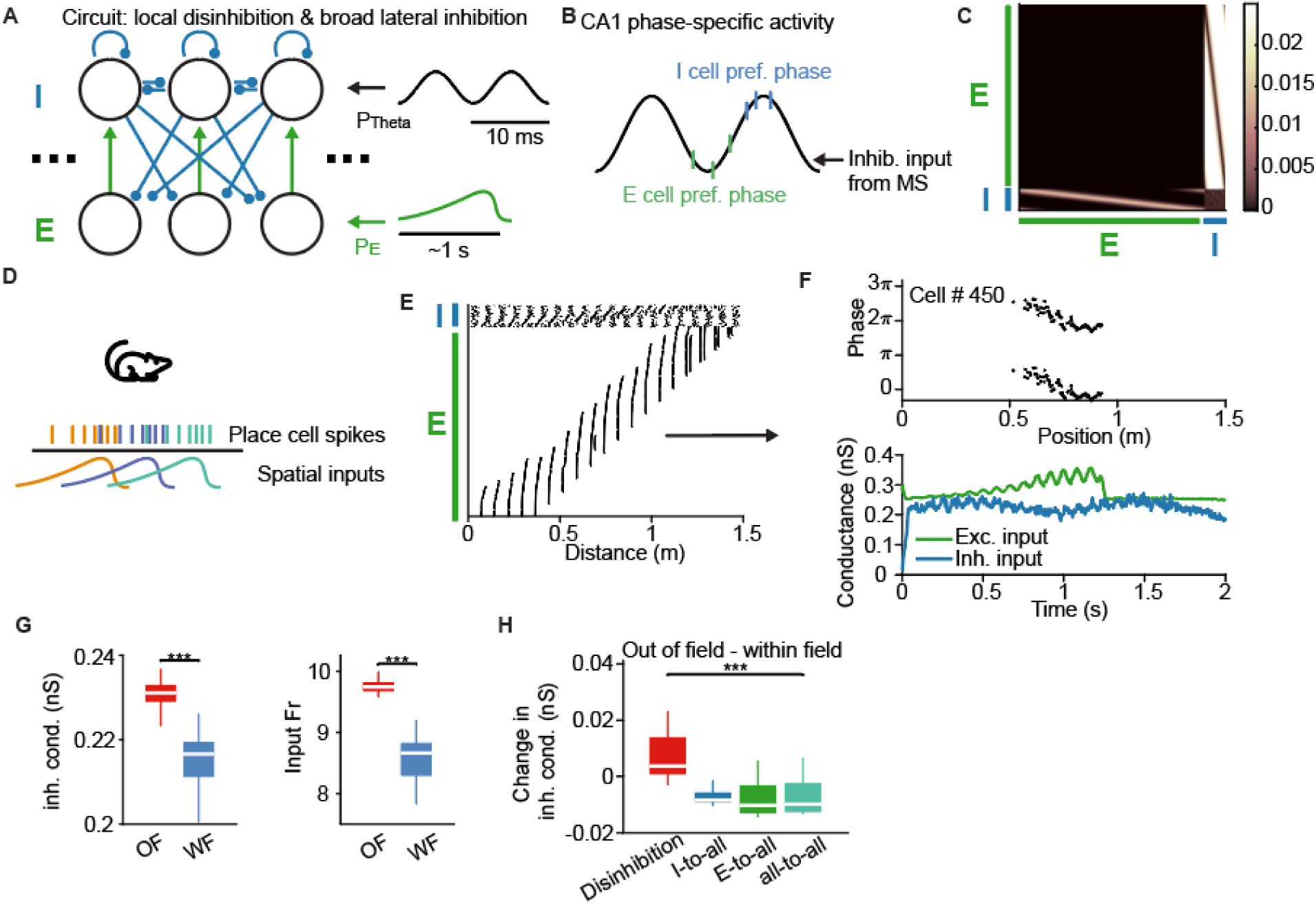
Spiking model illustrates sequential place field activity. **A:** Simplified circuit diagram of the spiking model with a disinhibitory motif (3 excitatory and inhibitory cells shown, out of a total of 1000 leaky integrate-and-fire cells **B:** Theta phase preference in CA1. Excitatory cells are active at the trough of the local field potential theta rhythm, while inhibitory cells tend to be active near the peak of theta. Theta input is modeled as inhibitory input from the medial septum to inhibitory cells. **C:** Weight matrices for outgoing excitatory and inhibitory connections. **D:** Schematic of simulated behavior and stimuli. **E:** Example of model behavior from one simulated trial. **F:** Phase-related activity of a single excitatory cell, averaged over 15 trials. Top: position vs. phase spiking activity of cell 450 along the track. Bottom: Excitatory (green) and inhibitory (blue) input conductance to cell 450. **G:** Mean inhibitory input within all model place fields. Left: Trial-averaged inhibitory conductance to excitatory cells outside of a place field (OF) versus within the place field (WF). Right: The firing rate of presynaptic inhibitory cells. White bar marks the data median, the rectangle marks the lower and upper quartiles, and the whiskers mark the data minimum and maximum. **H:** Comparison of inhibitory input to place cells outside of a place field versus within a place field for different connectivity schemes; inhibitory (I)-to-all, excitatory (E)-to-all, and all-to-all circuit connectivity.

We simulated an animal navigating along a linear track at 75 cm/s (4). Each excitatory cell received spatio-temporally organized input, simulating place-field-like input from CA3 (Fig. 1D). Spatial input was an increasing then decreasing ramp, as suggested in (38,41). There was an oscillatory component coupled to the input to mimic phase precession in CA3 place cells.

The model replicated place cell-like activity within the hippocampus. Excitatory cells exhibited theta oscillation-timed spiking activity within their place field and were silent otherwise (Fig. 1E). A cell’s activity ‘phase-precessed’ through the place field, spiking earlier on each theta cycle than during the previous cycle (Fig. 1E, F-upper). Inhibitory cells were locked to the theta phase, spiking reliably near/just prior to the peaks of the theta-timed input. An example place cell is shown in Figure 1F, demonstrating phase precession from the late to early phase of the theta oscillation. Inhibitory input conductance to the same cell decreased within the place field compared to out of the place field, consistent with recent findings that somatic inhibitory input to excitatory cells decreases within their place field (20,21; Fig. 1F-bottom). We ran 15 trials with uniformly distributed start phases of the theta oscillation. The average inhibitory input conductance, across all place fields and trials within the model, decreased significantly (p < 0.001, Kruskall-Wallis) within a place field compared to out of the place field (Fig. 1G). Similarly, the firing rates of the presynaptic inhibitory cells decreased significantly within the place field compared to outside the place field (p < 0.001, Kruskall-Wallis).

We tested several circuit configurations, adjusting coupling strength to keep the total inhibitory and excitatory coupling to a cell at a constant level (see Methods), so that we could compare changes in single-cell activity due to circuit connectivity alone (Fig. 1H). We did not observe the same decrease in inhibitory input without the disinhibitory motif (Fig. 1H). That is, when inhibitory cells had all-to-all excitatory connectivity to excitatory cells, excitatory cells had all-to-all inhibitory connectivity to inhibitory cells, or both, inhibitory input increased rather than decreased within the place field.

### Place field dynamics with disinhibition for an idealized 2-population model

We next created a minimal 2-population model of the disinhibitory circuit to understand the relation between the observed disinhibitory effect and connections between populations in the model. We recreated the local disinhibition with broad lateral inhibition circuit in two populations by modeling two excitatory populations (E_1_ and E_2_) that projected only to their local inhibitory population partners (I_1_ and I_2_ respectively, ‘on-center’) while the inhibitory populations projected only to the opposite excitatory cell population (‘off-center’, Fig. 2A). The inhibitory populations also inhibited themselves and each other. Excitatory populations received a ramping input while inhibitory populations receive a theta-timed oscillatory input.

**Figure 2.**
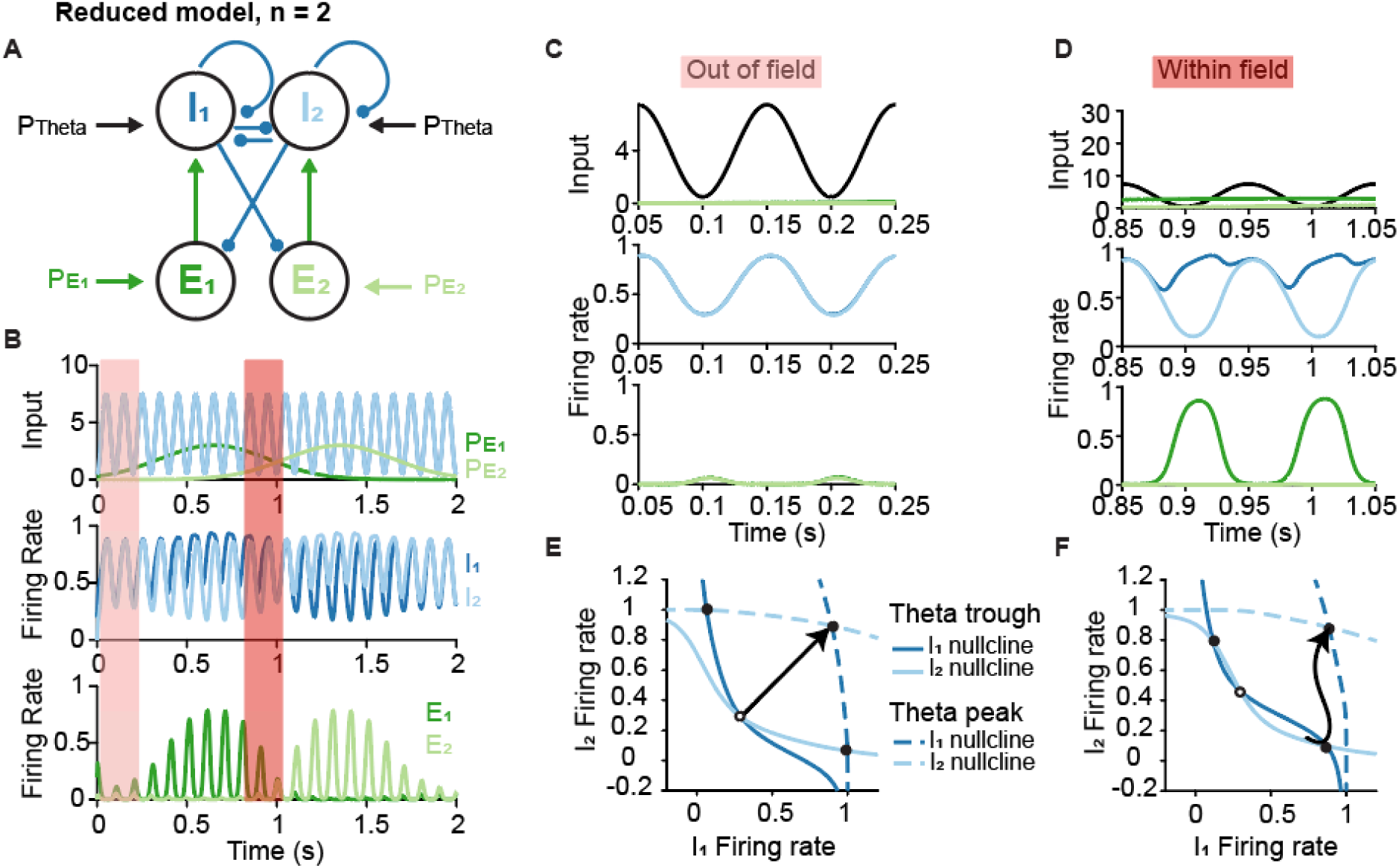
Equivalent rate model with disinhibitory motif shows qualitatively similar behavior to spiking model. **A**: Diagram of rate model. Excitatory cells received a depolarizing input within their place field (P_E1_ and P_E2_), while inhibitory cells received an ongoing theta-like oscillatory input (P_Theta_). **B**: Time course of model activity. Top row: inputs to excitatory populations are in green (dark green: input to E_1_, P_E1_; light green: input to E_2_, P_E2_); while an ongoing oscillatory input to the inhibitory populations, P_Theta_, is in black. Middle row: Firing rates of the inhibitory populations (Dark blue: I_1_; light blue: I_2_). Bottom row: Firing rates of the excitatory populations (Dark green: E_1_; light green: E_2_). Pink and red highlighted areas are shown in panels (C) and (D). **C**: Inset of the pink zone in (B). Top: Inputs P_E1_, P_E2_, and P_Theta_. Middle: Firing rates of I_1_ and I_2_. Bottom: Firing rates of E_1_ and E_2_. **D**: Inset of the red zone in (B). Layout is the same as in (C). **E**: Nullclines of I_1_ (dark blue) and I_2_ (light blue) corresponding to time points in (C). Solid lines correspond to time points when P_Theta_ is high, and dashed lines correspond to time points when P_Theta_ is low. Solid circles are stable fixed points, while hollow circles are unstable fixed points. **F**: Nullclines of I_1_ and I_2_, similar to (E) but corresponding to the time points in (D).

This minimal rate model reproduced key features of the spiking model. Specifically, the rate model generated place cell-like activity and allowed us to identify the conditions under which excitatory populations exhibit place field-like responses (Fig. 2B). When excitatory input was low for both excitatory populations, the excitatory populations remained silent while the inhibitory populations were active (Fig. 2C). This corresponded to the out-of-field (OF) case, where connections between populations are minimally effective in eliciting activity (Fig. 2C). Within the place fields (within-field, WF) a different dynamic emerged: one excitatory population (E_1_), exhibited strong firing within-field, while the other (E_2_), just entering its place field, had a very low firing rate (Fig. 2D). In this scenario, the inhibitory population, I_1_, connected to E_1_ became strongly active and suppressed E_2_. In contrast, I_2_, which did not receive strong excitatory drive, is less active and provided weaker inhibition to E_1_. This asymmetric inhibition produced a disinhibitory effect that supported selective excitation within the place field, illustrating how the model implemented a disinhibitory circuit motif.

### Inhibitory population model follows theta input outside of a place field

We further investigated the properties of disinhibition in the model by a geometric assessment of the model’s dynamics using phase plane visualization of the activity of the inhibitory populations. We observed biased competition between the inhibitory populations by examining the conditions for disinhibition when computing the inhibitory population nullclines. The I_1_ and I_2_ nullclines are the two curves in the firing rate plane along which dI_1_/dt and dI_2_/dt are zero, respectively. Intersections of the nullclines define fixed points (steady states), which can be either attractive (stable), or repulsive (unstable).

First, we focused on the Out of Field case. We found that when theta-timed input to the inhibitory cells is high, both inhibitory populations demonstrate stable, high firing rates (Fig. 2C, E, dashed lines). When theta-timed input to the inhibitory cells was low, we observe three fixed points (Fig. 2E, solid lines). This configuration is typical of competition: either one population is dominant over the other, corresponding to the two stable points; the ‘middle’ steady state is unstable – it is a saddle fixed point. When there was low input to the excitatory populations, the inhibitory populations both maintained an identical oscillation between a moderate and high firing rate (Fig. 2E, solid black arrow). While the middle state was unstable, the symmetry of the nullclines and the relatively fast 100 ms timescale of theta relative to the inhibitory population timescale, *τ_i_* = 10 ms, ensured similar I_1_ and I_2_ firing rates on each theta cycle. However, any additional noise in the model would have had the potential to bias activity on individual theta trials. Such noise-induced switching behavior in the model may be related to ‘flickering’ between place cell maps at decision points (42,43).

### Model inhibitory population demonstrates biased competition within a place field

Next, we examined the Within Field case. Within the place field of E_1_, the additional excitatory input from E_1_ altered the shape of the I_1_ nullcline and shifted it upwards (Fig. 2F). The result was that the I_1_ and I_2_ nullclines no longer intersected along the main diagonal, generating a larger basin of attraction towards the stable fixed point corresponding to high I_1_ and low I_2_ firing rates. The firing rate of I_1_ and I_2_ would then move towards this fixed point corresponding to a high I_1_ firing rate and low I_2_ firing rate on each theta cycle. This nullcline configuration corresponded to a biased competition between the inhibitory populations and demonstrated how the disinhibitory circuit displays pattern-generating activity. This disinhibitory pattern was necessary for generating the release from inhibition in both the idealized model and full spiking model (Fig. 2E, F), suggesting that the results from our idealized model generalize to the spiking model. Biased competition signifies a winner-take-all (WTA) behavior within a circuit when an excitatory population receives input. The tendency towards winner-take-all improves the signal-to-noise of place cell activity in the presence of input noise (44–46).

### Inhibitory-to-excitatory connections control disinhibitory behavior

We next determined the parameter range that sustained the competition state by performing a bifurcation analysis on the reduced model. To display the attractor/repeller features of the competition dynamics, we plotted the steady states as a function of input to the excitatory cells, P_E_ (Fig. 3A). At a low value of P_Theta_ and P_E_ = 0, there was a single stable state with a low firing rate of I_1_. Similarly, there was a single stable state at high values of P_E_, corresponding to a high firing rate of I_1_. At intermediate values of P_E_, there was a bistability in the firing rate of I_1_, the signature of WTA competition. This bistability was represented as the three possible I_1_ firing rates between PE values of approximately 2-6 in figure 3A, two stable, represented by solid lines, and one unstable, represented by a dashed line. The P_E_ values at the ‘knees’ defined the range of bistability and emergence/disappearance of the steady state coalescing with the saddle, a so-called saddle-node bifurcation.

**Figure 3.**
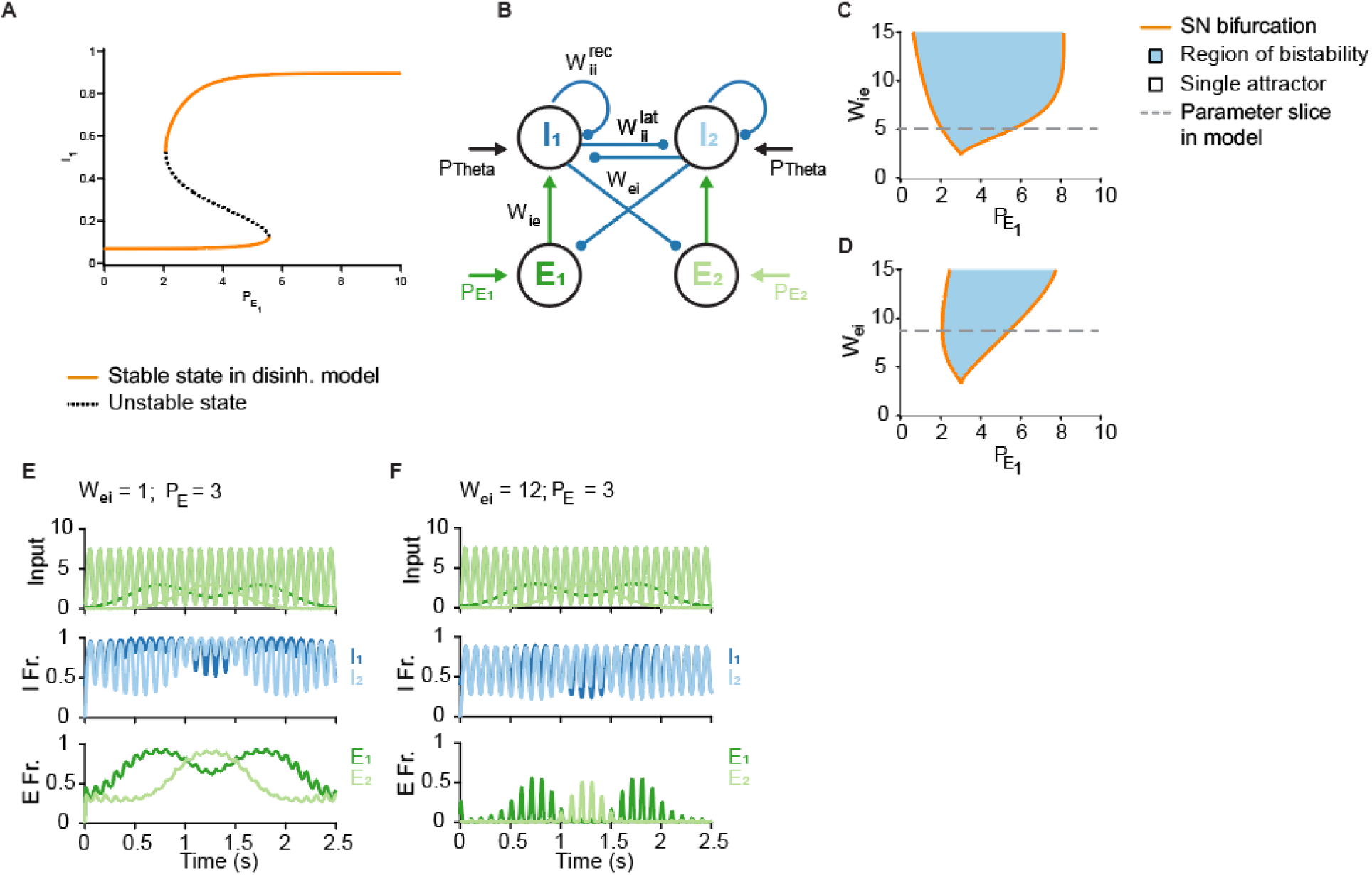
Input strength and inhibitory-to-excitatory weight important for control of disinhibitory effect. **A:** One-parameter bifurcation diagram of I_1_ firing rate in response to increasing input to excitatory cells. **B:** Diagram of reduced disinhibition model. **C:** Two-parameter bifurcation diagram for I_1_ state with respect to W_ie_ and P_E_. **D:** Two-parameter bifurcation diagram for I_1_ state with respect to W_ei_ and P_E_. **E:** Time course of model activity with reduced inhibitory-to-excitatory weight (W_ei_ = 5). Excitatory activity reflects the inputs, with high out-of-field activity and high within-field firing rates. **F:** Time course of model activity with increased inhibitory-to-excitatory weight (W_ei_ = 25). Excitatory activity is sparse and within-field firing rates are lower than in (E).

The saddle-node bifurcation could be computed directly (47) and they defined the boundaries of regions where bistability of competition would occur if we were to vary a second parameter, say W_ie_ (Fig. 3C). We could then determine the parameter regime at which a saddle-node bifurcation occurred. We found that a non-zero value of excitatory-to-inhibitory coupling, W_ie_, was required for the emergence of the competition state, but that otherwise, excitatory-to-inhibitory coupling did not exert much influence over the existence of competition within the model at a range of P_E_ (Fig. 3C). We found that the inhibitory-to-excitatory coupling, W_ei_, influenced the range of P_E_ values that sustained the competition state (Fig. 3D). The implication here was that altering W_ei_, but not W_ie_, could improve the winner-take-all properties of the network.

To demonstrate this effect, we examined two example scenarios: one with low inhibitory-to-excitatory coupling (Fig. 3E, W_ei_ = 5), and one with high inhibitory-to-excitatory coupling (Fig. 3F, W_ei_ = 25). With low W_ei_, the model was not in the region of bistability and the firing rates of the excitatory populations reflected the value of their inputs. Each population showed high firing rates both outside and within its place field. At a high W_ei_ value, the model was in the region of bistability and the excitatory populations had no activity outside of their place fields and sparser activity within the place field. This transition from broad to sparse tuning was a result of a change in inhibitory-to-excitatory weight alone, and mimicked the change in place field representation from novel to familiar environments (48–51).

### Disinhibitory control is more flexible without recurrent excitation

A hallmark of recurrent excitation is the trade-off between persistent activity thought to be important for memory (7,8,26) and the tendency towards runaway excitation (52). Disinhibition without recurrent excitation is a mechanism for maintaining persistent activity that is unlikely to lead to runaway excitation. We tested these concepts in our reduced 4-population model of disinhibition by illustrating the competition dynamics as analogous to a bistable system, with attractors viewed as minima in a double-well potential energy. Using long-duration simulations of the reduced model, we computed the dwell time in the disinhibition model subject to injected white noise (53). Mean dwell time is a measure of the height of the energy barrier when switching between two stable states, and as such is a metric of the rigidity of a model (Fig. 4A). A mean dwell time of zero corresponds to a lack of distinct stable states in the model, while a low (non-zero) dwell time corresponds to frequent switching, or a highly flexible model. A high (and non-infinite) mean dwell time corresponds to an inflexible model with discrete states. We found that increasing inhibitory-to-excitatory coupling in the model led to moderate mean dwell times in excitatory and inhibitory populations, while increasing recurrent excitation in the model resulted in mean dwell times approximately 1-2 orders of magnitude higher – increased rigidity (Fig. 4B,C). Increasing inhibitory-to-excitatory coupling therefore improved signal-to-noise in neural activity while maintaining flexibility (Figs. 3F, 4B). We found that even a small degree of recurrent excitation led to longer dwell times (Fig. 4C). The relation between recurrent excitation and sustained activity recapitulates work demonstrating that recurrent excitation coupled with disinhibition is involved in bursting activity in the recurrent excitiatory CA3 region (54,55), and that CA3, which is also a frequent generation site for epileptic activity (56).

**Figure 4.**
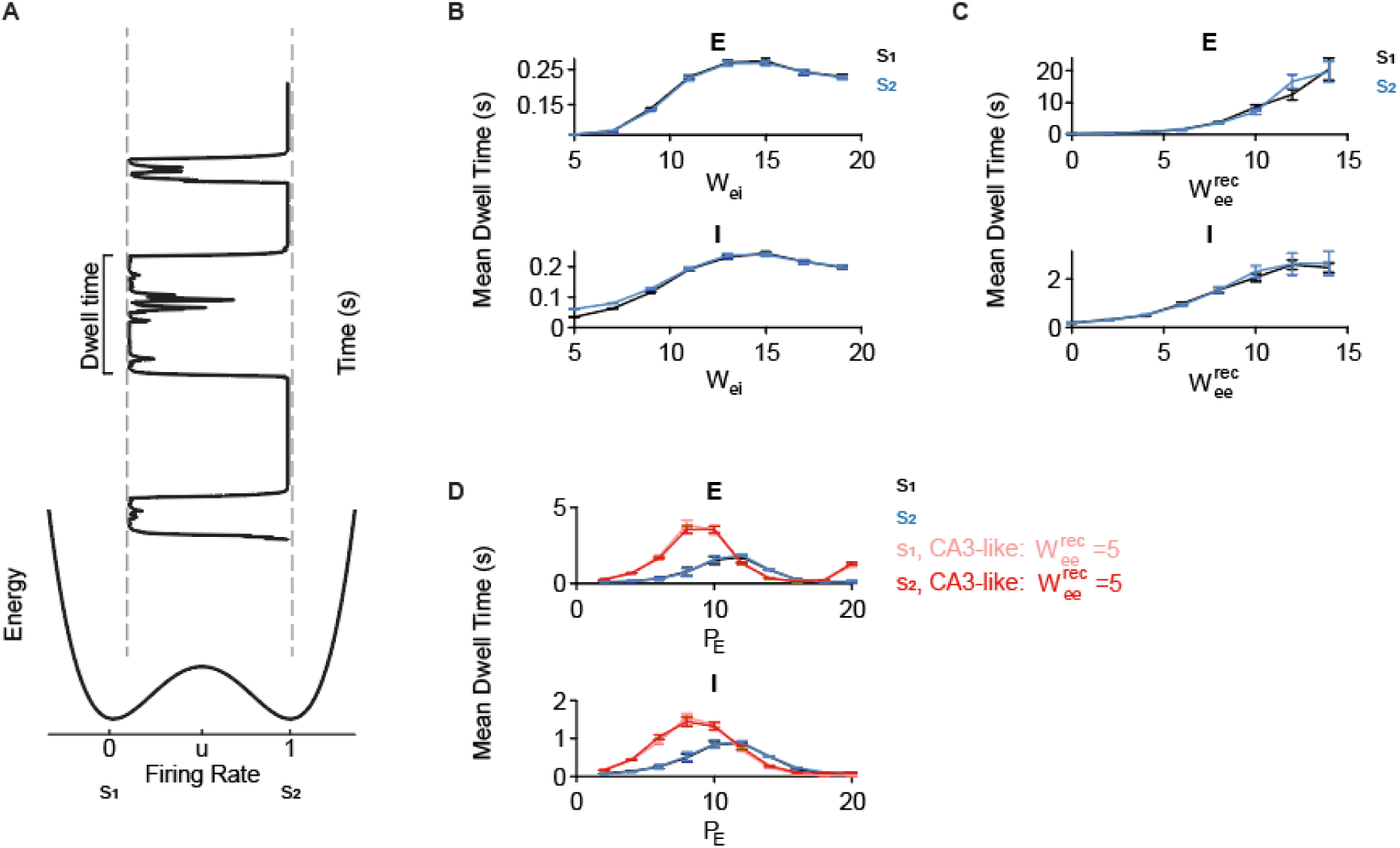
Adding recurrent excitation to the model reduces flexibility. **A:** Representation of the energy function of the firing rate of cells within our model. The points s1 and s2 correspond to the stable fixed points in 2H, while the point u corresponds to the unstable fixed point. Depending on the height of the barrier at u, it takes more (or less) energy to switch between s1 and s2, corresponding to longer (or shorter) dwell times in each state. **B:** Dwell times of excitatory (top) and inhibitory (bottom) cells with increasing inhibitory-to-excitatory weight (W_ei_). **C:** Dwell times of excitatory (top) and inhibitory (bottom) cells with increasing recurrent excitation (W_ee_^rec^). **D:** Dwell times of excitatory (top) and inhibitory (bottom) cells with increasing input to excitatory cells (P_E_), given the default disinhibition model (blue) versus a CA3-like disinhibition model with recurrent excitation (W^rec^ = 5).

We next compared our disinhibition model versus a CA3-like version of the model with *W^rec^* = 5 at increasing levels of P_E_ (Fig. 4D). We found that the original CA1-like disinhibition model demonstrated lower peak mean dwell times and accepted a wider input dynamic range, both traits that suggested greater flexibility in the CA1-like model versus the CA3-like model. The lower peak dwell times relate to the concept of faster remapping in CA1 versus CA3 (12), in addition to the idea of CA3 as an autoassociative network with stronger pattern separation in CA3 versus CA1 (26,57,58).

### Disinhibition motif allows for improved place field representation through increasing inhibitory-to-excitatory coupling

Noise correlations are the correlation between the activity of two neurons that is beyond shared input and instead due to circuit connectivity (59–61,61,62). They are typically quantified as the correlation in the fluctuations of individual trial responses between neurons for a given stimulus. For example, two neurons may be more active than their overall mean firing rate in response to the same spatial bin (assuming space is the input stimulus to place cells), and less active than their overall mean in response to another spatial bin, resulting in a positive noise correlation value between these two neurons. The opposite is also possible – one neuron can consistently increase its activity higher than average while another consistently decreases its firing rate in response to the same stimulus, leading to a negative noise correlation. In contrast, correlations that are purportedly due to shared input are called signal correlations and are calculated as the correlation between mean firing rates of pairs of neurons. When noise and signal correlations share the same sign, the information capacity of a network is reduced and the ability to distinguish between the responses of the two neurons is reduced as well, compared to the scenario where noise and signal correlations have opposite signs (63).

One way to measure improvements in learning is to measure the slope of the relationship between signal and noise correlations. A steeper slope signifies greater discriminability (64,65). We measured the signal-noise slope within our model by plotting noise correlations between neurons against the distance between place field centers for neuron pairs (Fig. 5). In the case of interneurons, which do not receive sharp spatially-modulated input, we instead examined the noise correlation relationship compared to the place field preference of their local excitatory cells.

**Figure 5.**
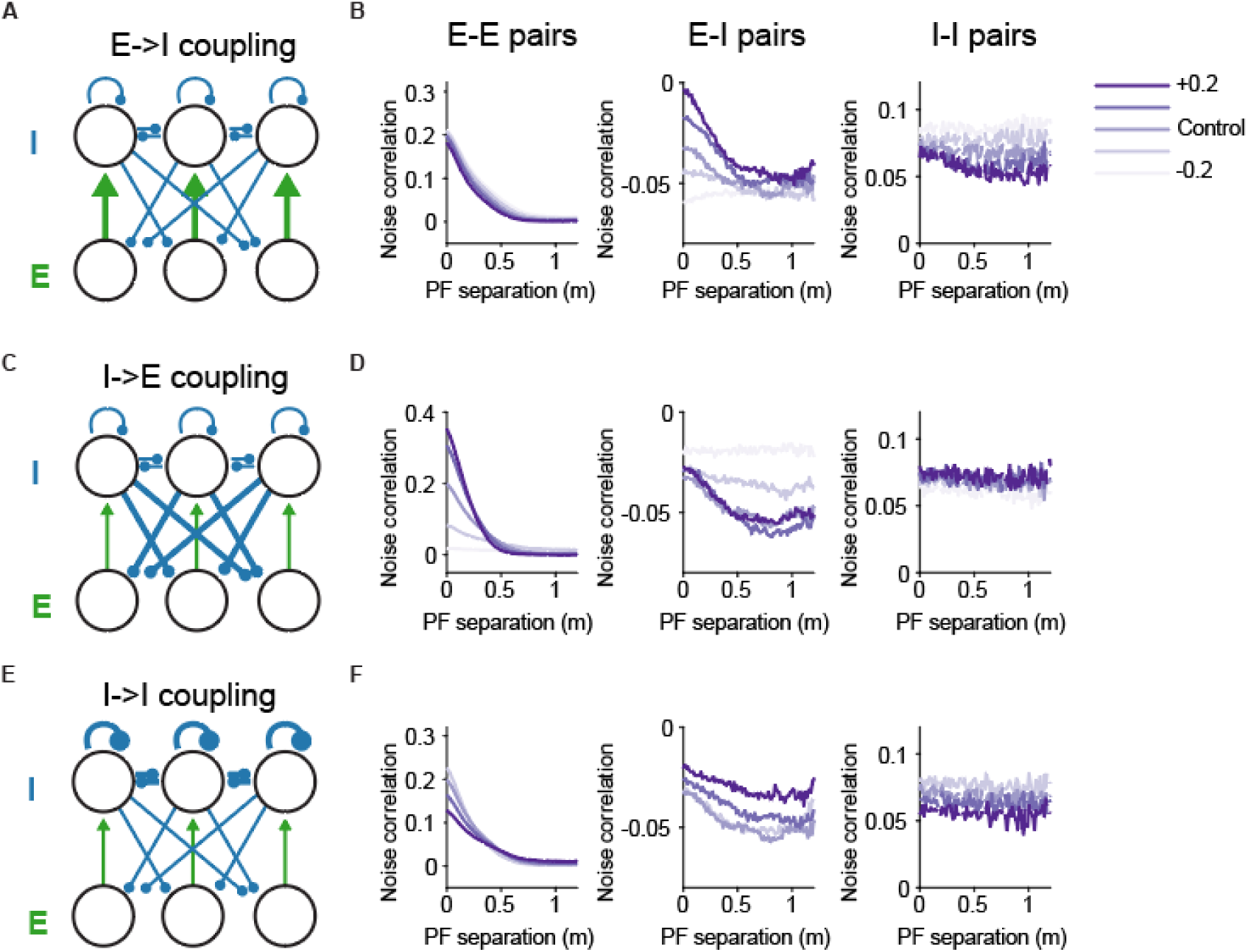
Effects of disinhibition circuit motif on representation. **A:** Schematic that illustrates increasing excitatory-to-inhibitory coupling. **B:** Excitatory-inhibitory cell pair noise correlations are most affected by altering excitatory-to-inhibitory coupling. Pairwise noise correlations compared to place field (PF) separation distance between excitatory pairs (left), excitatory-inhibitory pairs (middle), and inhibitory-inhibitory pairs (right). Inhibitory place fields were determined by the preferred place field location of the paired excitatory cells. Lighter purple denotes decreasing the excitatory-to-inhibitory weight by 0.1-0.2, while darker purple denotes increasing the weight by 0.1-0.2. **C-D**: similar format to A-B, for altering inhibitory-to-excitatory coupling. **E-F:** similar format to A-B and C-D, for altering inhibitory to inhibitory coupling.

Noise correlations are often used to determine ‘functional connectivity’ between cells. However, in highly recurrent networks, noise correlations as a metric of functional connectivity can lead to spurious connectivity relations (66). We demonstrate that disinhibition also generated predictable spurious correlations between excitatory cells. We ran the spiking model with independent white noise (mean of 0 and standard deviation of 0.2) injected into each LIF neuron. We then calculated the noise correlations by calculating the pairwise correlations between cell responses on each trial, after subtracting the mean response across 20 trials (see Methods for further detail). The signal correlations in this paradigm were the distance between place field centers.

Though there were no connections between excitatory cells in our model, we still observed positive noise correlations between nearby place cells, due to overlapping regions of disinhibition (Fig. 5B, D, F). These positive noise and signal correlations were consistent with the general finding that cells with similar stimulus encoding tend to have positive noise correlations (67–69).

We found that the only way to steepen the signal-noise slope was to increase the inhibitory-to-excitatory connections or decrease the inhibitory-inhibitory connections in the network (Fig. 5C, D, left). Altering the excitatory-to-inhibitory weights had no effect on the signal-noise slope of excitatory cell pairs. Surprisingly, the only way to alter noise correlations between inhibitory cells was to increase the coupling from excitatory to inhibitory cells.

### Disinhibition enables intrinsic sequential activity

We built a model that supports endogenous sequences based on the previous disinhibition model, adding feedforward disinhibition. As before, inhibitory populations had off-center coupling to excitatory populations, while excitatory populations had on-center coupling to inhibitory populations. We added excitatory coupling from each excitatory population to the next inhibitory population, which resulted in decreased inhibition to the next excitatory population, feedforward disinhibition (Fig. 6A, B).

**Figure 6.**
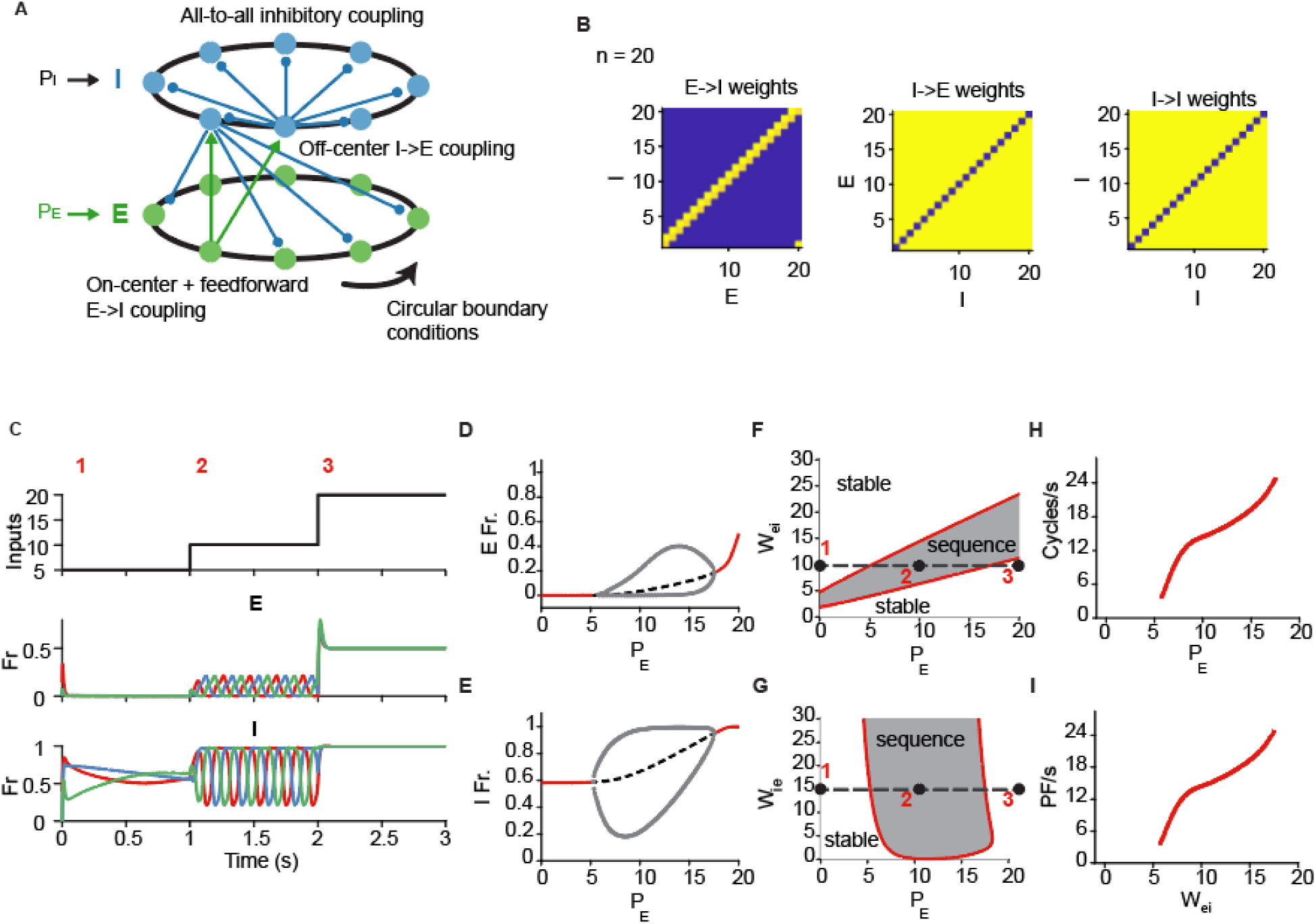
Disinhibition model of sequential activity. **A:** Schematic of model connectivity. Populations are arranged on a one-dimensional ring. Inhibitory populations exhibit all-to-all coupling, with no recurrent inhibition. Inhibitory populations connect to off-center excitatory populations and excitatory populations connect in an on-center manner with a feedforward connection to generate feedforward disinhibition. Inputs to the excitatory population (*P_E_*) and to the inhibitory population (*P_I_*) are uniform. **B:** Coupling matrices for the excitatory-to-inhibitory coupling (top left), inhibitory-to-excitatory coupling (top right), and inhibitory-to-inhibitory coupling (bottom right). Connections are binary, instead of following a Gaussian footprint. **C:** Example activity within the model with n = 3. Top: inputs to the excitatory population. Middle: activity of each excitatory population. Bottom: Activity of each inhibitory population. **D:** One-parameter bifurcation diagram for an excitatory population in response to increased input *P_E_*. **E:** One-parameter bifurcation diagram for an inhibitory population in response to input to the excitatory population. **F:** Two-parameter bifurcation diagram demonstrating the region of the Hopf bifurcation in the *W_ie_* and *P_E_* parameter space. Black dots denote the values of *W_ei_* and *P_E_* in the time course in (A). **G:** Two-parameter bifurcation diagram demonstrating the region of the Hopf bifurcation in the *W_ie_* and *P_E_* parameter space. As in (F), the black dots denote the values of *W_ie_* and *P_E_* in the timecourse in (A). **H, I:** Relationship between the frequency of the sequential activity and *P_E_*.

The model displays both winner-take-all and sequential activity dynamics, in a manner that depends on input to the excitatory and inhibitory populations (Fig. 6C). We used a ring architecture for convenience; a chain without a ring structure can also sustain sequential activity.

Our goal was to understand how the level of tonic (and uniform across units) excitatory drive controls behavior of the model, simulating the changes in excitatory drive when the entorhinal cortex and CA3 are silenced. We first analyzed an idealized 3-population version of the model (Fig. 6C), though we subsequently demonstrated that model behavior is invariant to the number of populations in the model. The firing rate of the excitatory population underwent a Hopf bifurcation at intermediate levels of *P_E_* – the emergence of sequential activity in this 3-population model. The inhibitory population also underwent a Hopf bifurcation in the parameter range (Fig. 6E). Yet instead of a population-level synchronous oscillation, we observed a temporally periodic traveling wave pattern in the excitatory and inhibitory populations, with all units oscillating in sequence. The sequence functioned as a moving WTA profile, mirroring the tendency towards winner-take-all in networks with lateral inhibition without sequential activity (70–72).

We then extended our analysis to find the region of the Hopf bifurcation in both the range of *W_ei_* and *P_E_*. We found a monotonic relationship between the existence of the Hopf bifurcation and increasing values of Wei and *P_E_* (Fig. 6F). This region approximately corresponded to the range of values where −*W_ei_I* + *P_E_* (from the above equation for the activity of the excitatory population) was above some small threshold.

We found a different pattern for *W_ie_*’s impact on periodic activity in the model. Increasing *W_ie_* had little effect on the existence of the Hopf bifurcation or its relationship with *P_E_*. We also found that increasing *W_ei_* and *P_E_* increased the frequency of the sequence in a non-linear manner (Fig. 6H, I).

## Discussion

We have investigated the computational significance of disinhibitory circuits in the hippocampal CA1 region in both spiking models and idealized rate models. Disinhibition accounts for the decrease in inhibitory input to place cells when spiking within a place field. Additionally, disinhibitory circuits provide a mechanism for increasing inhibitory-to-excitatory coupling to enhance place field selectivity consistent with experimental literature.

Our model offers an explanation for recent experimental results demonstrating that presynaptic inhibition decreases within a place cell (20,21). The model can sustain low firing rate sequential activity even when excitatory input to the model is low, as demonstrated in recent experimental findings (16). The model is invariant to the number of neuron populations in the network, which is important since there are hundreds to thousands of neurons in the mammalian hippocampus (73,74). We also found that we could regulate the behavior of the model by the inhibitory-to-excitatory neuron coupling strength and drive to the excitatory population.

### Role of interneurons in control of CA1 pyramidal cells

Prior reports disagree on a potential role inhibition in turning pyramidal cell activity. Some works suggest a non-differential (“blanket inhibition”, 75) effect in tuning, whereas others attribute a key role in shaping pyramidal neurons’ firing (20–22,37), although these experiments tested only tested one type of interneuron at a time.

We propose that PV+ interneurons, inhibited by SST+ interneurons could match the activity profile suggested by a disinhibition model of circuit activity and warrants further exploration (22,76–78). Interestingly, in the study by Chamberland, Nebet, et al. (2023), only PV cells that received weak excitation exhibited a decrease in firing rate. This observation, together with the fact that hippocampal interneurons exhibit non-overlapping connections onto pyramidal cells (79), offers an anatomical basis for our disinhibitory circuit model. Depolarization-induced suppression of inhibition due to retrograde endocannabinoid signaling at CCK+ synapses onto pyramidal cells may also be involved in the reduction in inhibitory input throughout a place field (80–82). Although the precise interneuronal connectivity needs to be identified, disinhibition is a common motif in the brain to enhance excitatory activity (83–86). Elucidating the potential role of disinhibition and the patterns generated in large circuits is important for understanding complex responses of in vivo circuits.

### Computational implications of attractor-like models on rate-remapping

We explored the role of biased competition between two attractors, simulating the interactions between place cells in a circuit. Previous models underemphasize the role of local circuitry and largely neglect ongoing activity in hippocampal circuits. We used a novel switching circuit to demonstrate several desired general properties of place field networks, including sparse activity due to competition, decreasing firing rates with familiarity, and temporally changing positive noise correlations at short time scales. The theta-phase-specific, attractor-like activity in our model has implications for place-field dynamics that remain underexplored. The self-regulating property of attractor networks suggests a mechanism for scaling place field size with the environment (87,88). Further, our model may explain findings that place fields are more expansive in large environments (89,90) and smaller and more numerous in visual cue-rich environments (91,92). The effects of lateral inhibition may be weaker when place fields are few and larger, allowing more flexibility in place fields, and stronger when place fields are numerous, forcing smaller place fields.

### Theta period-timing of disinhibitory activity

We used our model to gain insight into ‘flickering’ behavior observed when animals exhibit goal-directed behavior within a maze, and place cell assemblies alternate activity between theta cycles (42,43,93,94). The distribution of cycle occupancy towards representation of one environment versus the other on each theta cycle may reflect the relative strength of inputs towards the respective assembly of place cells.

The theta-timed disinhibitory attractor characteristic in our model also implies that the greatest changes in firing rate will occur within a confined phase span of the theta cycle. Decreases in presynaptic inhibitory neuron firing rate were strongest at the trough of theta, away from the imposed preferred theta phase of the interneurons in our simplified model (Fig. 2G). As a result, it is difficult to detect strong changes in firing rate when averaging over several trials in vivo when theta phase does not align across trials (95). Instead, increases in phase-specificity coupled with a decrease in firing rate may be necessary to detect changes in presynaptic activity.

### Implications and extensions of the sequential activity model

Our one-dimensional model of homogenous populations generated sequences that repeated in the same order. While this is different from in vivo findings that the hippocampus generates sequences of varying lengths involving different, non-repeating cells (96), it was previously shown that a model of sequence generation (with recurrent excitation in CA3) could generate multiple different sequences through a toroidal connectivity that acts as an analogous extension to two dimensions of the one-dimensional ring connectivity in our model (29). Thus, a two-dimensional version should be able to generate sequences of varying length involving different, non-repeating cells as well. Further, since we demonstrated that sequential activity depends on the coupling from inhibitory to excitatory populations, heterogeneously-coupled populations may demonstrate sequences of varying lengths that reflect a random draw of population order within a sequence. When the sequential activity arrives at an inhibitory-excitatory pair where the coupling strength between the inhibitory population and the next excitatory population is too high, the sequence would end. This sensitivity to input strength and inhibitory-to-excitatory gating may also reflect the increased likelihood of participation of high-firing rate pyramidal cells in ripples and place cell activity (54,88,97). These high-firing rate cells may have connectivity that places them closer to a disinhibitory regime than low-firing rate cells. Sequences could also end through facilitating plasticity from excitatory populations to a tertiary population of slow-activating interneurons. Our goal with the current model was to determine how an idealized disinhibition-based model of sequential activity can be used to provide a framework for understanding sequential activity in CA1 circuits.

### Potential sources of sequential connectivity

Our model is highly structured and assumes forward connections between excitatory populations to inhibitory populations to attain sequential activity. The sequential structure may be preconfigured during development (98,99). Alternatively, it could be formed through plasticity mechanisms combined with repeated offline sequential activity during sleep or rest (100–102) as an animal familiarizes itself with an environment (27,103).

### Limitations of population models

Hippocampal activity is high dimensional, relative to motor cortex or sensory cortex, for example, and exhibits a hyperbolic geometry, suggesting highly complex circuitry and dynamics (104–107). Our reduced model is deliberately idealized to two populations of excitatory and inhibitory cells, omitting biophysical morphology of cell types and the wide variety of interneuron subtypes. Nonetheless, we still observe biased competition and changes in E-I balance within our model at a level of detail compatible with modern calcium imaging or electrophysiological recording techniques. Potential future modeling directions would be to test disinhibitory circuits in a small number of two-compartment models and to test the effects of multiple interneuron types, many of which target dendrites instead of the soma (21,22,108,109).

## Materials and Methods

### Spiking model description

The spiking model consisted of 894 excitatory neurons and 904 inhibitory interneurons. Each neuron was modeled as a conductance-based integrate-and-fire neuron. The change in membrane potential, *V_i_*, of the *i^t^*^ℎ^ neuron is given by:

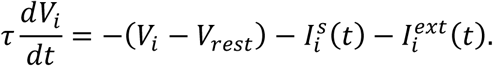

The value *V_rest_* is the resting membrane potential, 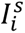 is the synaptic input current, and 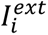 is the external input current. The neuron ‘spikes’ once *V_i_* reaches threshold, *V_t_*_ℎ*res*ℎ_, and is manually reset to *V_rest_*. Parameter values are outline below.

The synaptic input is the sum of excitatory (*I^ex^*) and inhibitory (*I^in^*) inputs:

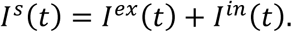

Synaptic input currents are defined by:

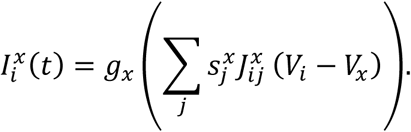

Here, *g_x_* is the conductance, 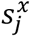 is the synaptic gating variable, *J^x^* is the connectivity matrix, and *V_x_* is the reversal potential, where *x* denotes either excitatory or inhibitory values. The gating variable, *s*, is defined by:

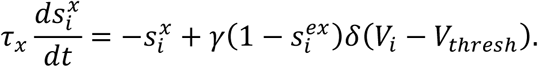

The connectivity matrix is defined by:

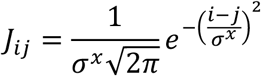

for the on-center connections, and

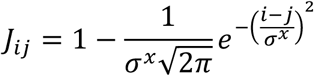

for the off-center connections, where *σ^x^* is the standard deviation of the width of the on-center or off-center connection.

The external input to the place fields is defined by a constant component with amplitude *I*_0_ and a skewed gaussian, centered on the place field start location, *x_i_*:

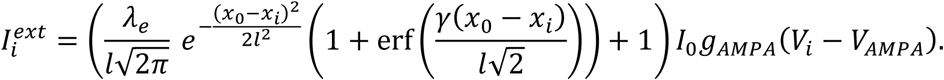

The parameter *λ_e_* defines the input amplitude, the parameter *l* defines the input width, *x*_0_ defines the location of the animal, and *γ* defines the skew of the input.

The oscillatory input to the inhibitory neurons was modelled as constant component with amplitude *I*_0+0.1_ and a sinusoidal component with amplitude *λ_i_*, period *T*, and phase offset *φ*.

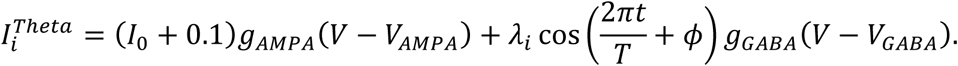

Place field centers are spaced uniformly along the track from 0m to 1.5m.

The model was solved numerically with Euler’s method and a timestep of 0.1ms. White noise with mean 0 and standard deviation of 0.2 was added to the simulation using the Orstein-Uhlenbeck process (110) and simulated with the Euler-Maruyama method with a timestep of 0.1 ms with the equation:

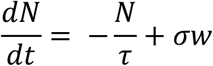

where *τ* = 10, *σ* = 0.2, and *w* is a white noise process. Parameters were chosen by adjusting parameters given in (27), then running a grid search through J_ei_, J_ie_, and J_ii_ that maximizes the decrease in inhibitory input within the place field, using NYU Langone’s High Performance Computing Core. All computations were done using Matlab 2019b.

**Table 1.** Spiking model parameters.

| Model Parameter | Description | Value |
| --- | --- | --- |
| $\tau$ | Membrane time constant | 20 ms |
| $V_{rest}$ | Resting membrane potential | -65 mV |
| $V_{thresh}$ | Action potential threshold | -50 mV |
| $g_{AMPA}$ | AMPA conductance | 0.2 pS |
| $g_{GABA}$ | GABA conductance | 2 pS |
| $V_{AMPA}$ | AMPA reversal potential | 0 mV |
| $V_{GABA}$ | GABA reversal potential | -85 mV |
| $\tau_{ex}$ | Excitatory synapse time constant | 4 ms |
| $\tau_{in}$ | Inhibitory synapse time constant | 8 ms |
| $l$ | Input length constant | 0.3 |
| $\lambda_e$ | Excitatory input amplitude | 0.16 |
| $\gamma$ | Input skew | 0.4 |
| $I_0$ | Constant input amplitude | 1.499 |
| $\lambda_i$ | Inhibitory input amplitude | 0.16 |
| $T$ | Period of oscillatory input | 100 ms |
| $J_{ie}$ | Synaptic strength from E to I | 0.015 |
| $J_{ei}$ | Synaptic strength from I to E | 0.02 |
| $J_{ii}$ | Synaptic strength from I to I | 0.02 |
| $\sigma^{IE}$ | Standard deviation of off-center connection from inhibitory to excitatory populations | 12 |
| $\sigma^{EI}$ | Standard deviation of the on-center connection from excitatory to inhibitory populations | 0.8 |

### Rate model description

Each population was modelled as a Wilson-Cowan-like equation. For excitatory populations, the excitatory population firing rate, *E*, was modelled as:

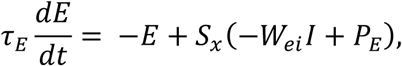

where *τ_E_* is the excitatory population time constant, *W_ei_* is the coupling strength from inhibitory to excitatory populations, *I* is the firing rate of presynaptic inhibitory populations, and *P_E_* is the external input to the population. Similarly, the inhibitory population firing rates were modelled as:

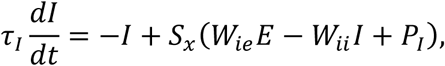

where *τ_i_* is the inhibitory population time constant, *W_ie_* is the coupling strength from excitatory to inhibitory populations, *W_ii_* is the coupling strength from inhibitory to inhibitory populations, and *P_i_* is the external input to inhibitory populations.

The function *S_x_* is the Boltzmann function:

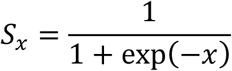

The input to the excitatory population was modelled as a skewed gaussian (38,41):

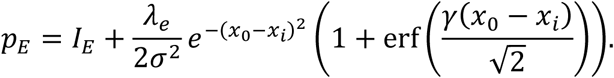

The input to the inhibitory population was modelled as a sinusoidal input:

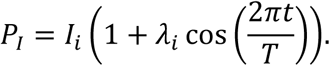

The model was solved numerically with Euler’s method and a timestep of 0.1ms.

**Table 2.**
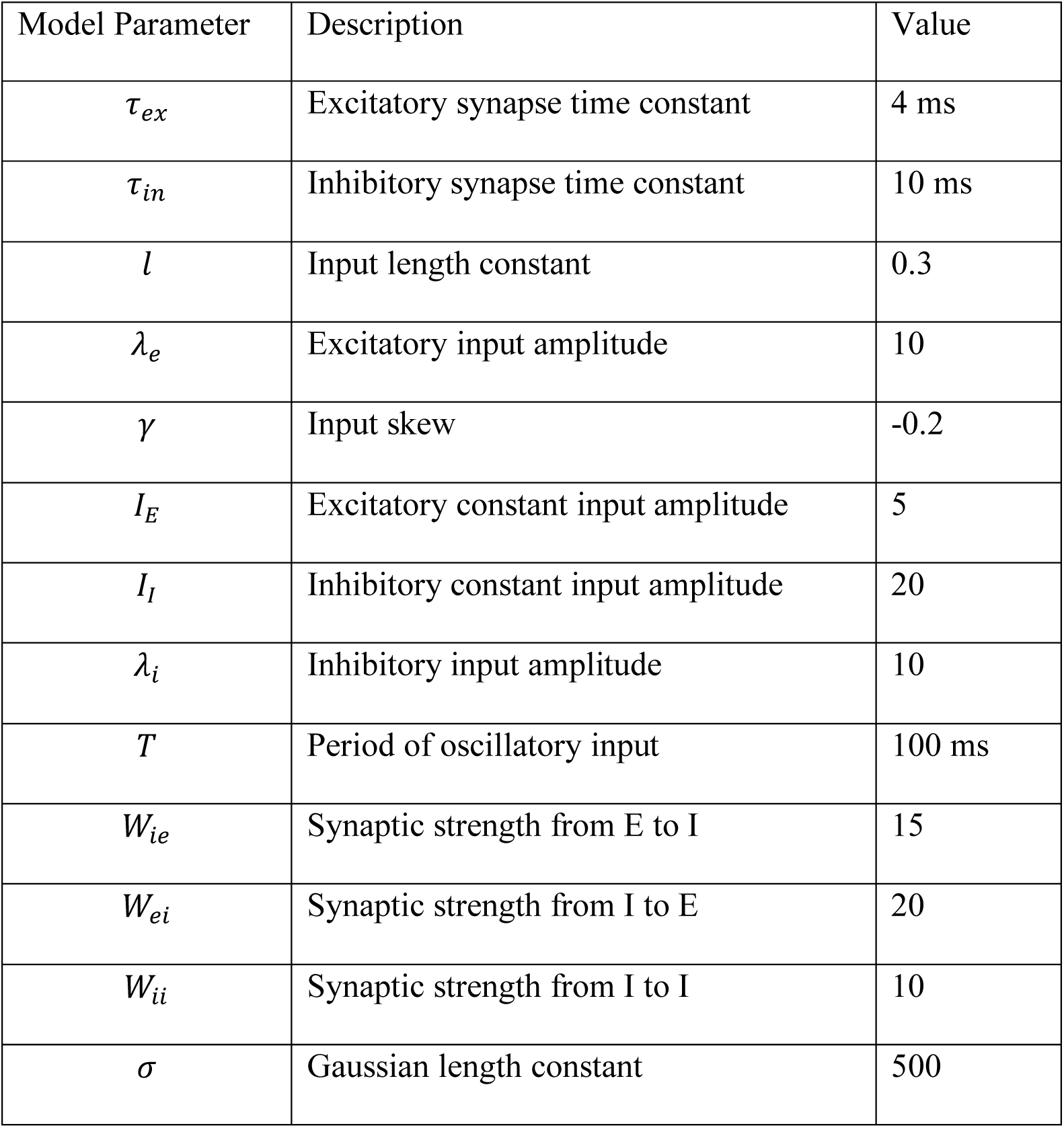
Rate model parameters.

**Table 3.** Rate model parameters, sequence model.

| Model Parameter | Description | Value |
| --- | --- | --- |
| $\tau_{ex}$ | Excitatory population time constant | 10 ms |
| $\tau_{in}$ | Inhibitory population time constant | 10 ms |
| $W_{ie}$ | Synaptic strength from E to I | 15 (except when noted) |
| $W_{ei}$ | Synaptic strength from I to E | 10 (except when noted) |
| $W_{ii}$ | Synaptic strength from I to I | 4 (except when noted) |

### Dwell time simulation

Excitatory cells in the rate model were injected with P_E_ = 3 plus white noise through the Orstein-Uhlenbeck process (110) and simulated with the Euler-Maruyama method with a timestep of 0.1 ms with the equation:

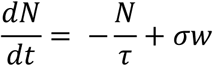

where *τ* = 10, *σ* = 2, and *w* is a white noise process. Each simulation was run for a model time of 100 s. Dwell time in each state was measured as the time spent in-between crossing a firing rate threshold of 0.5 Hz.

### Spiking model parameter calculation

Place field width was calculated as the width of the region within 10% of the maximum firing rate of the place cell. Maximum firing rate was calculated as the maximum value of the trial-averaged firing map.

Noise correlations were calculated as:

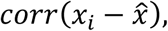

where *x_i_* is the binned spiking activity within each position bin on a single trial and *x̄* is the trial-averaged spiking activity. Spikes were binned in 2 cm bins.

#### Bifurcation Analysis

Bifurcation analysis was completed using Auto and XPP software (47).

## Lead contact

John Rinzel

## Data and code availability

All code and data will be made available upon publication.

## Acknowledgements

We thank members of the Rinzel and Buzsáki labs for feedback on the manuscript. This work was supported by R01 MH122391, 1R01MH139216. L.G. was supported by an NIH Training Grant for Computational Neuroscience (R90DA043849).

## Author Contributions

L.G., G.B., and J.R. designed the study. L.G. performed the computational modelling. M.V. collected data. M.V. and L.G. performed analysis of the data. L.G. and J.R. wrote the paper. All authors contributed to manuscript editing.

## Declaration of Interests

The authors declare no competing interests.

## Notes

### Competing Interest Statement

The authors have declared no competing interest.

